# Intraspecific interactions in the annual legume *Medicago minima* are shaped by both genetic variation for competitive ability and reduced competition among kin

**DOI:** 10.1101/695908

**Authors:** Sara Tomiolo, Claire Thomas, Michael K. Jespersen, Christian F. Damgaard, Bodil K. Ehlers

## Abstract

Knowing which mechanisms drive the outcome of intraspecific interactions is highly relevant for understanding diversity maintenance. Plant species that exhibit strong genetic substructure over small spatial scales may be exposed to frequent interactions with closely related individuals. Predictions of how genetic similarity may drive the outcome of intraspecific interactions are based on two contrasting theories: the resource partitioning hypothesis and kin selection theory. The first predicts that competition will be stronger among closely related conspecific (i.e. kin) because similar genotypes have similar resource requirements. The second predicts instead that competition will be reduced among kin, in order to maximize the inclusive fitness. Although efforts have been made to reconcile these two theories as non-mutually exclusive, the outcomes of intraspecific interaction studies are frequently interpreted as the results of either one or the other. We experimentally tested the hypothesis that intraspecific interactions may be driven by both genetic variation for competitive ability and reduced competition among kin. We used an annual legume, *Medicago minima*, to conduct two greenhouse experiments testing changes in root behaviour, above-ground growth and biomass in response to neighbour identity. We found evidence of both genetic variation for competitive ability and reduced competition among kin in some genotypes. Reduced competitive growth towards kin was found in the most competitive genotypes, suggesting that kin avoidance and competitive ability were simultaneously affecting plant behaviour and growth. With presence of both kin competition avoidance and variation for competitive ability, the outcome of intraspecific interactions will strongly depend on the local spatial genetic substructure. This is highly relevant to predict how intraspecific competition affect diversity maintenance.

## Introduction

The last decade has seen an increasing interest in understanding the mechanisms driving the outcome of intraspecific interactions and whether new insights can be integrated into species coexistence theory (e.g. Bolnick et al., 2011; Hart, Schreiber, & Levine, 2016). In plants, genetic variation for competitive ability for traits like growth rate, resource acquisition and allocation can create variation in competitive ability within a species. This often results in asymmetric competition where competitively superior genotypes outperform their neighbours (Weiner & Thomas, 1986; Weiner, Campbell, Pino, & Echarte, 2009).

Studies investigating the effects of genetic similarity on the outcome of intraspecific interactions generally test either of two theoretical predictions: the resource partitioning hypothesis (Young, 1981) and kin selection theory (Hamilton, 1964). The resource partitioning hypothesis, predicts that closely related species have similar resource requirements and therefore compete more harshly (MacArthur & Levins, 1967; Burns & Strauss, 2012). At the intraspecific level, where genetically similar individuals are expected to have similar resource requirements, the resource-partitioning hypothesis predicts that competition between close relatives should be stronger than between distantly related conspecifics (Young, 1981; Kelly, 1996). Conversely, kin selection theory predicts that kin should avoid strong competition with relatives, as it reduced the inclusive fitness (Hamilton, 1964) and should either reduce competitive interference with kin (West, Pen, & Griffin, 2002) or disperse to avoid competition (Ronce, 2007).

In plant ecology, resource-partitioning has long been the traditional view on plant competition. However, over the past decade kin selection theory has received support from a number of studies, which found that plants exhibit a plastic growth response with reduced competitive growth toward kin neighbours compared to unrelated conspecifics (e.g. Bhatt, Khandelwal, & Dudley, 2010; Dudley & File, 2007; Crepy & Casal, 2014).

There is currently no consensus, as empirical studies have produced contradicting results, finding evidence for both increased and reduced competition in plants growing with kin compared to non-kin (Donohue, 2003; Cheplick & Kane, 2004; Milla, Forero, Escudero, & Iriondo, 2009; Masclaux et al., 2010; Simonsen, Chow, & Stinchcombe, 2014). This is consistent with the view that only under a specific set of conditions would the evolution of reduced competition among kin be advantageous (Crepy & Casal, 2014; Ehlers & Bilde, 2019). For example, life history traits such as barochory and synaptospermy are associated with a clustered distribution of kin or siblings (Lepik, Abakumova, Zobel, & Semchenko, 2012), and in this case it would be favourable to discriminate between siblings and strangers, and modulate the intensity of competition accordingly.

One issue is that it can be difficult to determine if different outcomes of intraspecific interactions between kin and non-kin are caused by “actual” kin responses or by genetic similarity. With genetic variation for competitive ability, symmetric competition among kin may be the result of similar competitive ability among genetically identical individuals, but may also result from reduced competition toward plants recognized as kin. These two explanations can lead to similar outcomes in terms of mean fitness differences between kin and non-kin interactions, making it difficult to distinguish between the explanatory mechanisms (e.g. Milla et al., 2009; Simonsen et al., 2014). One way to disentangle these effects would be to monitor whether differences in performance among individuals change when plants are prevented from recognizing the identity of their neighbours. This can be attained by “silencing” root exudates that mediate neighbour recognition (Chen, During, & Anten, 2012). If within-group variation among individuals is determined exclusively by genotype-driven differences in phenotype, we expect similar variation in performance when plants can or cannot recognize their neighbours. However, if within group variation in plant performance is affected also by kin recognition, we expect within-group variation in performance to change when plants are prevented from recognizing neighbour identity.

Recent conceptual papers propose that, rather than interpreting intraspecific interactions as the results of either genetic variation in competitive abilities or avoidance of kin competition, both mechanisms may act simultaneously within a species and possibly even within a population (File, Murphy, & Dudley, 2011; Ehlers & Bilde, 2019). Namely, the same genotype could experience reduced intraspecific competition when growing close to a kin, and varying intensity of intraspecific competition when growing with non-kin, depending on differences in competitive ability and resource requirements among genotypes However, this interpretation of intraspecific interactions has so far received little empirical attention (but see Biernaskie, 2010). If both mechanisms co-occur, the outcome of intraspecific interactions in natural populations may depend greatly on small-scale population genetic structure. Understanding the different mechanisms that drive genotype-dependent interactions among conspecifics, is relevant to explain maintenance of intraspecific genetic variation. Moreover, if intensity of intraspecific competition vary with the genetic relatedness of competing plants, this can alter the relative strength of within *vs.* between species competition and thus affect predictions for species coexistence (Chesson, 2000; Vellend & Geber, 2005; Laird & Schamp, 2006; Bodil K Ehlers et al., 2016).

In this study, we used the annual legume *Medicago minima* to investigate the outcome of intraspecific interactions, by growing different genotypes with either kin or non-kin neighbours. We report the results of two experiments in which we tested for a) different responses in root growth toward kin and non-kin neighbours, and b) different responses in above-ground growth and biomass of plants grown in mini-communities with neighbours of varying relatedness.

We used these two experiments to test the following hypotheses: 1) *M. minima* genotypes reduce competition among kin by decreasing root growth towards kin *vs.* non-kin neighbours, and 2) the outcome of competitive interactions among genotypes is affected by both genotype-specific variation in competitive ability among non-kin and reduced competition with kin.

## Material and methods

### Study species

*Medicago minima* L. is an annual legume widespread in the Mediterranean basin. This species is mostly autogamous and produces spiny dehiscent fruits typically containing 5-8 seeds each. Fruits mostly disperse close to the maternal individual, as they remain on the withered mother plant until the end of life cycle, but they can also be passively dispersed by grazers when attaching to their fur and possibly by ants (Wolff & Debussche, 1999). Previous studies on a closely related annual species, *Medicago truncatula*, which has a similar reproductive and dispersal strategy as *M. minima* and shares its same habitat, found that *M. truncatula* is characterized by a well-defined population structure in which densely populated kin patches alternate with more heterogenous matrix of different genotypes (Bonnin, Ronfort, Wozniak, & Olivieri, 2001; Siol, Bonnin, Olivieri, Prosperi, & Ronfort, 2007). Due to the strong similarity in life-history traits between the two species, we expect that they also exhibit a similar population structure.

For the current experiment, four genotypes of *M. minima* were sampled at four different sites (one genotype per site) in the garrigue vegetation, in southern France. Seeds from one maternal plant per site were grown in the greenhouse for one generation to remove maternal effects. Because M. minima is autogamous, seedlings emerging from the same seed pod are full sibs, and hereon we refer to them as kin. Distance between sampling sites varied between 500 m and 2 km. Our sampling design allowed - in the absence of genetic data - to exclude that the genotypes chosen for this study were close relatives. The closeness of sampling sites assured that genotypes were from the same climatic conditions, however we cannot exclude that differences in microsite characteristics may have led to small-scale local adaptation of single genotypes. The same genotypes used for this study were previously found to have higher survival in kin groups than non-kin groups, indicative of reduced kin competition (Bodil K. Ehlers et al., 2016).

### Experimental set up

We conducted two greenhouse experiments to test the effects of neighbour plant relatedness (i.e. kin or non-kin) on below- and aboveground growth of *Medicago minima* plants. In both experiments, plants were grown in a mix of sand, greenhouse soil and vermiculite in proportions 1:1:1. To ensure germination, seeds were scarified with sand paper and placed on wet filter paper for 48 hours before sowing onto seed trays placed in a greenhouse. Greenhouse temperature was set to a minimum temperature of 15 C degrees. After germination, seedlings were transplanted into pots, and care was taken to match similar sized seedlings for the same pot.

#### Experiment 1

We set up a pairwise comparison experiment in which each of the four target genotypes was grown with either a kin or a non-kin neighbour. To account for the effect that genetic variation in competitive ability may have on the outcome of genotype-by-genotype interactions, we combined each genotype with two different non-kin genotypes, thus resulting in one kin treatment and two non-kin treatments for each focal genotype (Table 1). Due to limitation in seed availability, we were not able to set up a full factorial experiment. Each of these genotype combinations was replicated three times.

**Table 1:** combinations of kin (K) and non-kin (NK) treatments used for both experiment 1 and experiment 2. Additional kin and non-kin treatments where activated carbon was added, are marked with a “c”. The top row indicates the focal genotypes, while the column indicates the identity of the neighbour or surrounding genotypes

|  | Genotype focal | 102 | 114 | 121 | 130 |
| --- | --- | --- | --- | --- | --- |
| **Neighbours** | **102** | K + c | NK + c |  | NK |
|  | **114** | NK | K | NK |  |
|  | **121** | NK |  | K | NK |
|  | **130** |  | NK | NK + c | K +c |

Plants’ pairs were sowed at a distance of 2.5 cm in transparent plastic containers (30 cm x 25 cm x 3 cm) that allowed to observe and measure the direction of root growth. We kept the plastic containers tilted at a 50° angle, thus forcing roots to grow against the downward-facing surface. To avoid exposure of roots to light, we kept the downward-facing side of each container covered with a black plastic slate that could be removed when recording root growth. Growth and direction of roots were monitored twice a week. A transparent millimetre plastic foil glued over the container’s surface, allowed for accurately measuring root growth. Seedlings of *Medicago minima* have one primary root growing vertically, and a number of lateral roots developing subsequently from the primary root. Secondary roots can develop in any direction, but due to the experimental set up in which we inclined containers and reduced the width of the container, we forced lateral roots to grow in two dimensions, either toward or away from neighbours. This allowed easy categorization of roots growing either away or toward the neighbour plants (Appendix 1, Figure A1, A2)

After 6 weeks, when the roots of several plants reached the bottom of the containers we harvested above- and belowground biomass of each individual. Roots were carefully separated from soil, washed, oven dried at 70° C degrees for 72 hours. Above- and belowground biomass were weighed separately on a high precision scale.

After completion of the experiment, the millimetre paper foils on which root development was recorded, were scanned and processed using the imaging software Image-J (Lobet, Pagès, & Draye, 2011) to estimate root length.

#### Experiment 2

In this experiment we sowed four individuals of *M. minima* in 20 cm diameter pots (volume 4l). These pots were considered adequate to represent the plant density observed in natural conditions (Poorter, Bühler, Dusschoten, Climent, & Postma, 2012). One focal genotype was placed at the centre of the pot and was surrounded by three equidistant individuals. Focal genotypes were exposed to three different treatments based on surrounding plants’ identity: 1) *kin*, where the surrounding individuals belonged to the same genotype as the focal; 2) *mix*, where the surrounding individuals belonged to three different genotypes (i.e. surrounding genotypes were different from each other and from the focal); 3) *non-kin*, where the three surrounding individuals belonged to the same genotype, but were different from the focal. The three treatments represent different scenarios of relatedness among conspecifics, in which a genotype is surrounded by either three identical genotypes or by three stranger genotypes identical to each other, or by three different stranger genotypes.

Each of the four genotypes was exposed to kin, mix and two different combinations of non-kin treatments. The genotype combinations used for non-kin treatments were the same used for experiment 1 (Table 1). Treatments were replicated four times for each genotype (N pots = 64). Additionally, we exposed two selected kin and two non-kin treatments (three replicates for each) to application of activated carbon, by adding 20 ml activated carbon/l of soil (N pots = 12). Activated carbon is known to dampen the effects of root exudates (Callaway & Aschehoug, 2000; Callaway, Ridenour, Laboski, Weir, & Vivanco, 2005), which mediate neighbour identity recognition (Semchenko, Saar, & Lepik, 2014). Thus, by comparing pots treated with activated carbon and pots that were untreated, we aimed at testing whether plants modify their growth when unable to recognize the identity of their neighbours.

We recorded diameter and number of leaves of focal plants weekly to estimate early growth rate. For the first three weeks, we measured diameter for both focal and surrounding plants, and afterwards we restricted measures only to focal plants. Due to technical difficulties, we were able to count number of leaves on focal and surrounding plants only for the first three weeks. After 16 weeks, aboveground biomass of all plants (focal and surrounding) was harvested individually, dried at 70° C degrees for 72 hours and weighed using a high-precision scale (Mettled Toledo AX504, d = 0.1mg)

### Statistical analyses

#### Experiment 1

For each focal plant, we estimated root behaviour of focal plants by calculating the difference between the number of lateral roots growing away from and growing towards the neighbour, standardized by the plant’s total number of roots. The same calculation was applied to root length, which was measured in cm. We applied linear models to test the effect of focal genotype, treatment (kin *vs.* non-kin) and their interaction on root behaviour (creating separate models for number of roots and roots length). We carried out power analysis to estimate the power of the statistical tests given the significance threshold and the sample size of our experiment.

Subsequently, we tested the effects of focal genotype identity and treatment (kin *vs.* non-kin) on above- and below-ground biomass (using generalized linear models with gamma distribution and identity link function) and on the roots-to-total biomass ratio (using linear models). Identity of neighbour genotypes was not specified in the model due to the limited number of replicates available.

#### Experiment 2

We calculated early radial growth and early leaves growth as the difference in respectively maximum diameter and number of leaves between week 2 and week 1. We applied linear regression models to test how early leaves growth of focal and surrounding individuals varied across treatments. We also applied linear models to test for the effects of focal genotype, treatment (kin, non-kin and mix) and their interaction on early radial growth and final biomass of focal plants.

To test how total pot biomass and within-pot variance in biomass changed across treatments, we used linear models, and applied log-transformation to both total biomass and biomass variance. We also used linear models to test how total pot biomass and within-pot variance in biomass changed across treatments (kin *vs.* non-kin) in response to activated carbon. For this analysis, total biomass was square-root transformed and within-pot variance was log-transformed. Effects of focal genotype, treatment and surrounding genotype identity on focal plant’s biomass were tested using linear models.

All statistical analyses were conducted using the statistical software R version 3.5.3 (R Core Team, 2019). Post-hoc tests were carried out using pairwise comparisons in the package emmeans (Lenth, 2019). Statistical code for data preparation, statistical analyses and data visualization is documented in Appendix A2.

## Results

### Experiment 1

We found that root growth (both number and length of roots) away *vs.* toward a neighbour varied across focal genotypes and treatments (Table 2, Figure 1A). Two genotypes (114 and 121) grew more roots away from their neighbour when this was a kin compared to non-kin, a response consistent with avoiding root competition with kin. In contrast, genotype 102 grew more roots toward kin neighbours compared to non-kin, whereas no sign of a differential root growth in response to neighbour identity was observed for the fourth genotype (130). These results were consistent for number of roots (Table 2, Figure 1A) and for the length of roots (Table 2, Figure 1B). Power analysis indicated that the power of the test for number of roots was 0.83 and for root length 0.87.

**Table 2:** Results of linear models conducted on the number and the total length of roots growing away *vs.* toward a neighbour, in response to identity of focal genotype, treatment (kin or non-kin) and their interaction (Experiment 1)

| term | df | F.test | p.value |
| --- | --- | --- | --- |
| genotype_focal | 3 | 4.42 | 0.012 |
| treatment | 1 | 13.25 | 0.001 |
| genotype_focal:treatment | 3 | 7.68 | 0.001 |
| term | df | F.test | p.value |
| genotype_focal | 3 | 3.4 | 0.031 |
| treatment | 1 | 14.6 | 0.001 |
| genotype_focal:treatment | 3 | 8.37 | <0.001 |

**Figure 1:**
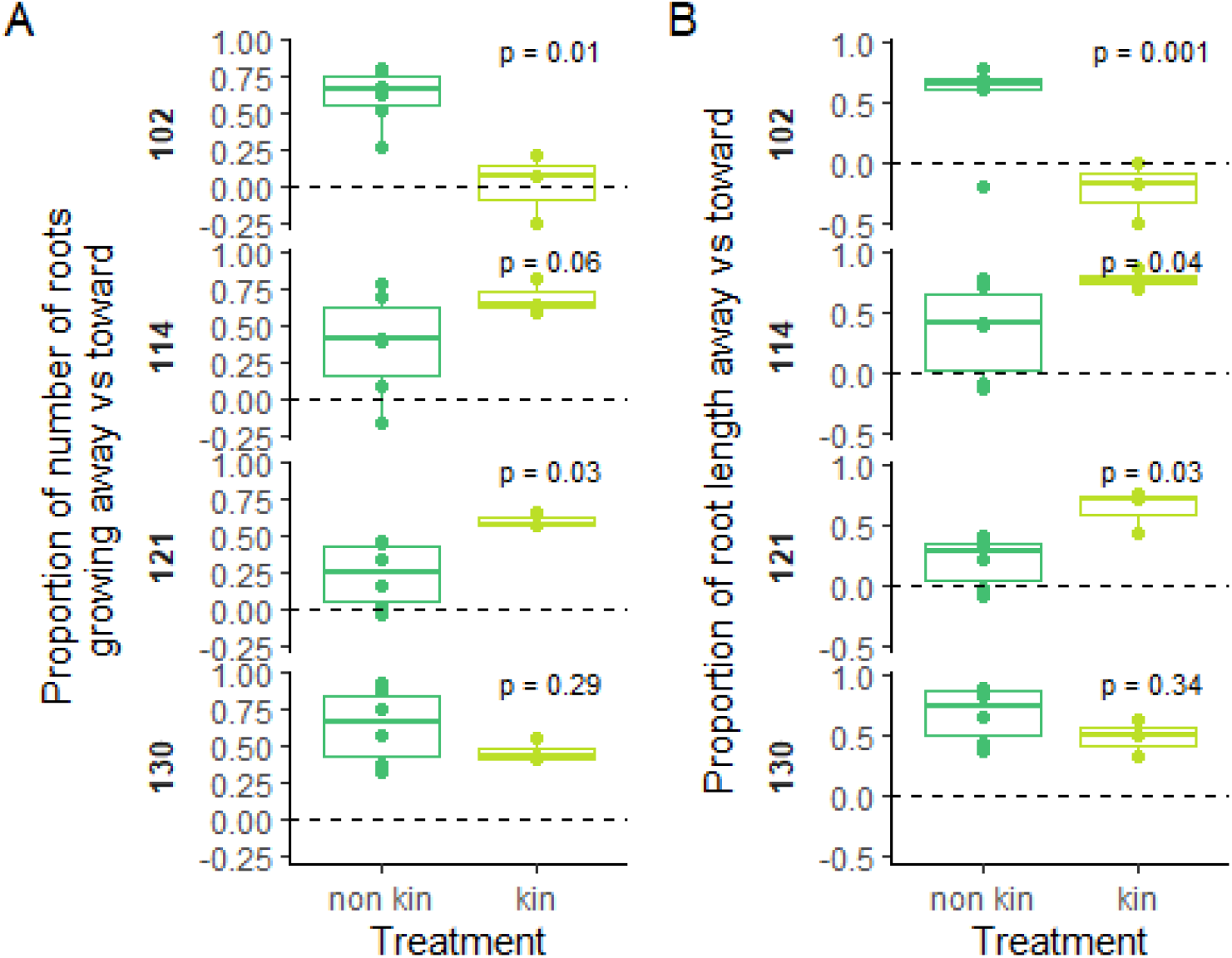
Boxplots of A) Relative number of roots growing away *vs.* toward the neighbour, B)Relative length of roots growing away *vs.* toward the neighbour (experiment 1). Each dot represents one individual replicate. Positive values indicate a higher number of roots and higher root growth (measured in cm) away than toward the neighbour. Panels represent focal genotypes (102, 114, 121, 130) separately, and on each panel we report the results of post-hoc tests for the effect of treatment (kin *vs*. non-kin).

We did not find any effect of treatment (kin *vs.* non-kin) on the production of aboveground biomass (Wald-Chi-Square = 1.39, p = 0.23), belowground biomass (Wald-Chi-Square = 0.35, p = 0.55) or on root-to-total biomass ratio (F_1_ = 0.14, p = 0.70). Identity of focal genotype had an effect only on belowground biomass (Wald-Chi-Square = 14.40, p = 0.002) where focal genotypes 102 and 130 produced the lowest amount of root biomass (Figure 2).

**Figure 2:**
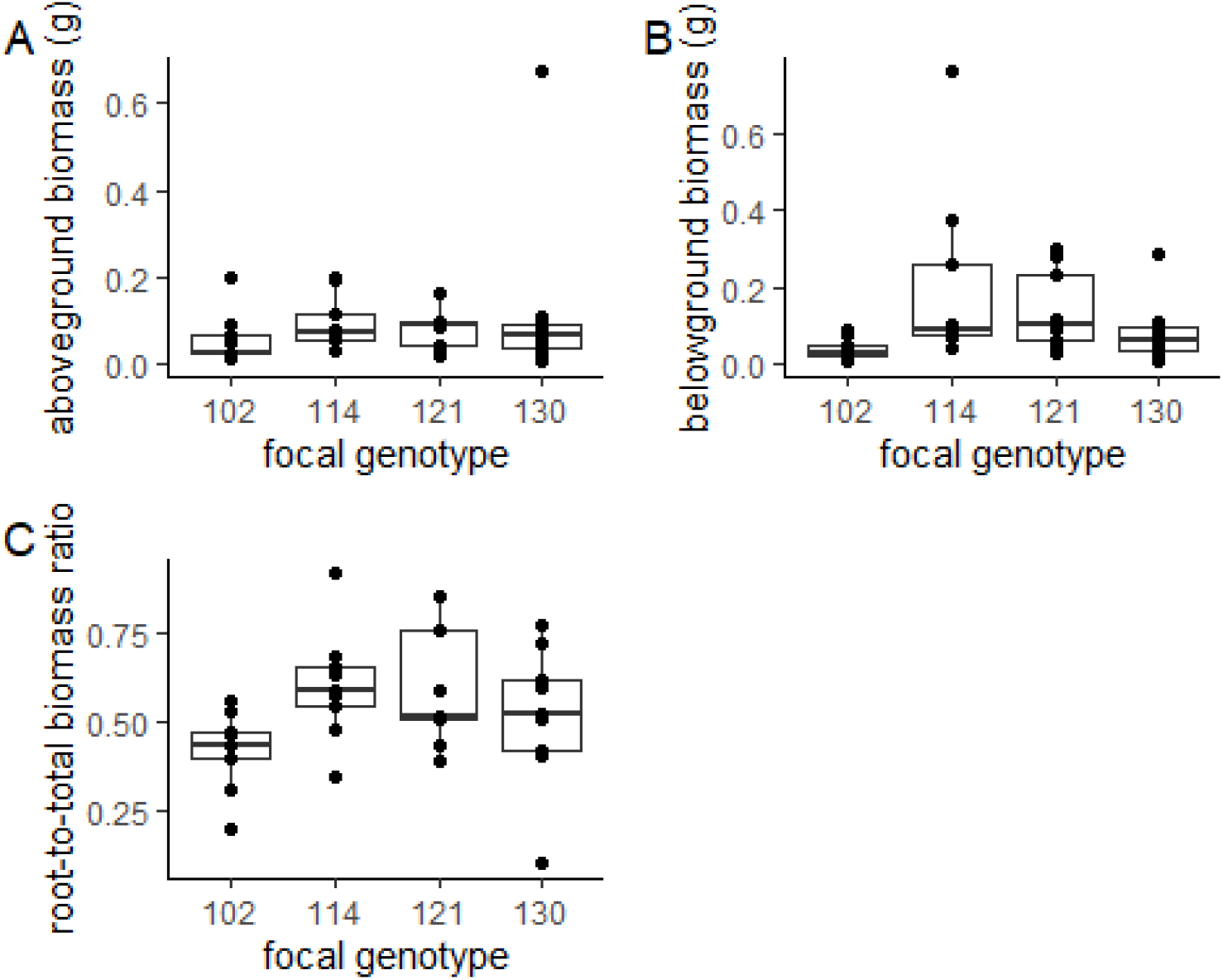
Boxplots of A) aboveground biomass, B) belowground biomass, C) root-to-total biomass ratio for the focal genotypes (experiment 1). Dots represent single focal plants.

### Experiment 2

Linear regression for growth of focal and surrounding leaves (Figure 3A-C) revealed different slopes across treatments. We found a significant positive relationship between leaves growth of focal and surrounding plants in kin treatments (F_1_=8.94, p = 0.009), whereas we did not find any significant relationship in mix (F_1_= 0.19, p = 0.64) and non-kin (F_1_= 0.15, p = 0.69) treatments. Early radial growth was different across genotypes (Table 3, Figure 3C). Genotype 102 attained the lowest growth rate and genotype 121 the fastest growth rate, but no significant effect of treatment was found (Table 3).

**Table 3:**
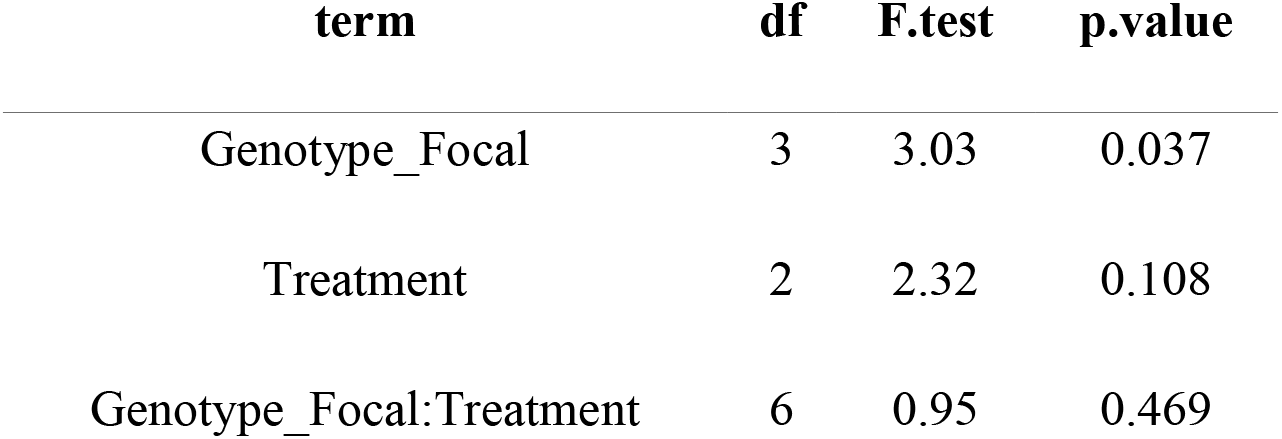
results of linear models for early radial growth of focal individuals in response to focal genotype and treatment (Experiment 2)

**Figure 3:**
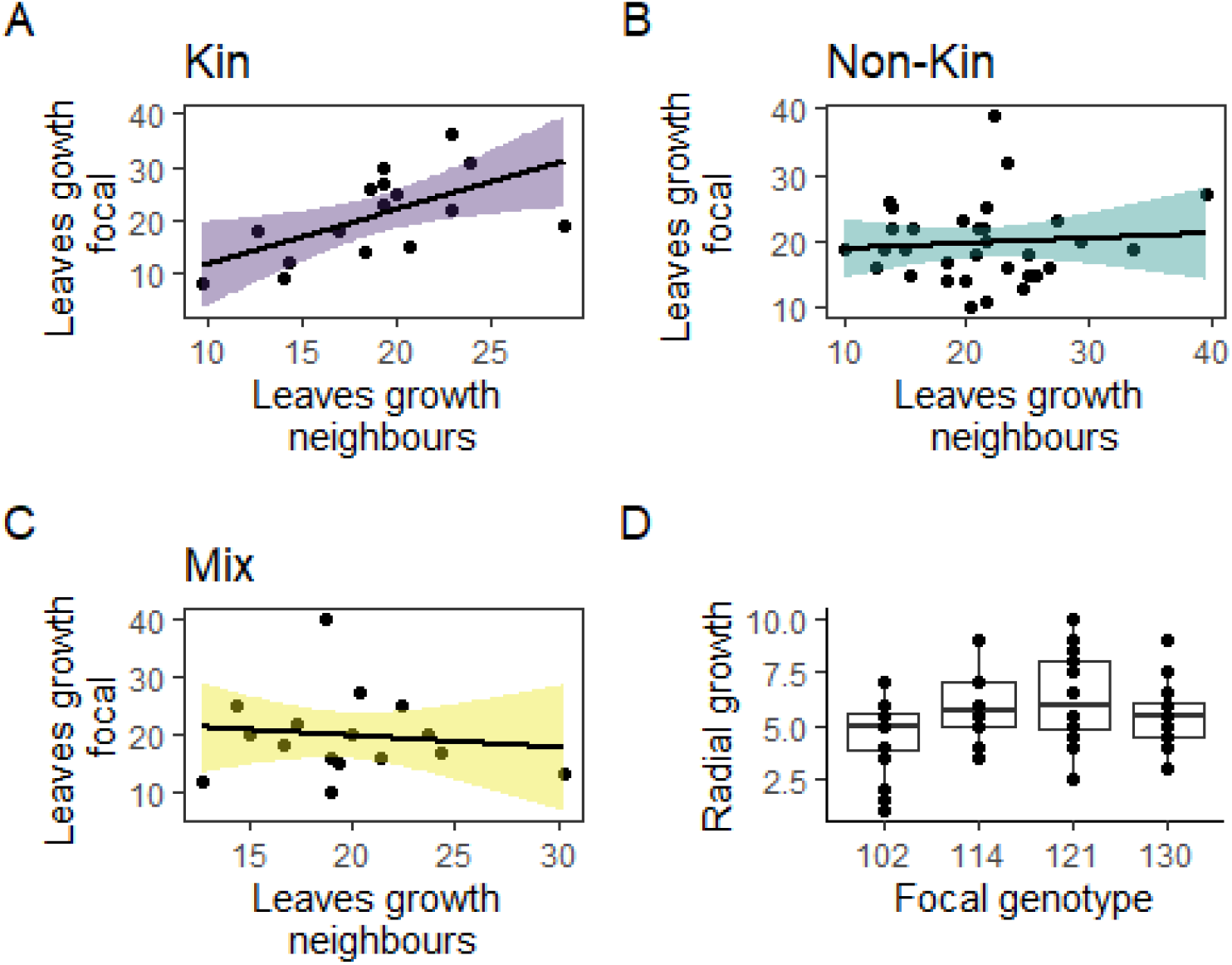
Regression for number of leaves grown during the first two weeks (experiment 2) on focal and surrounding genotypes across treatments (kin, non-kin and mix).

Analysing pot-level aboveground biomass, we found that total pot biomass (Figure 4A) was very similar across treatments (F_2_ =0.26, p = 0.7), in contrast the within-pot variance in biomass (Figure 4B) varied substantially across treatments (F_2_ =3.59, p = 0.03). The variance in biomass among plants within pots in the mix treatments was three times higher than in kin treatments.

**Figure 4:**
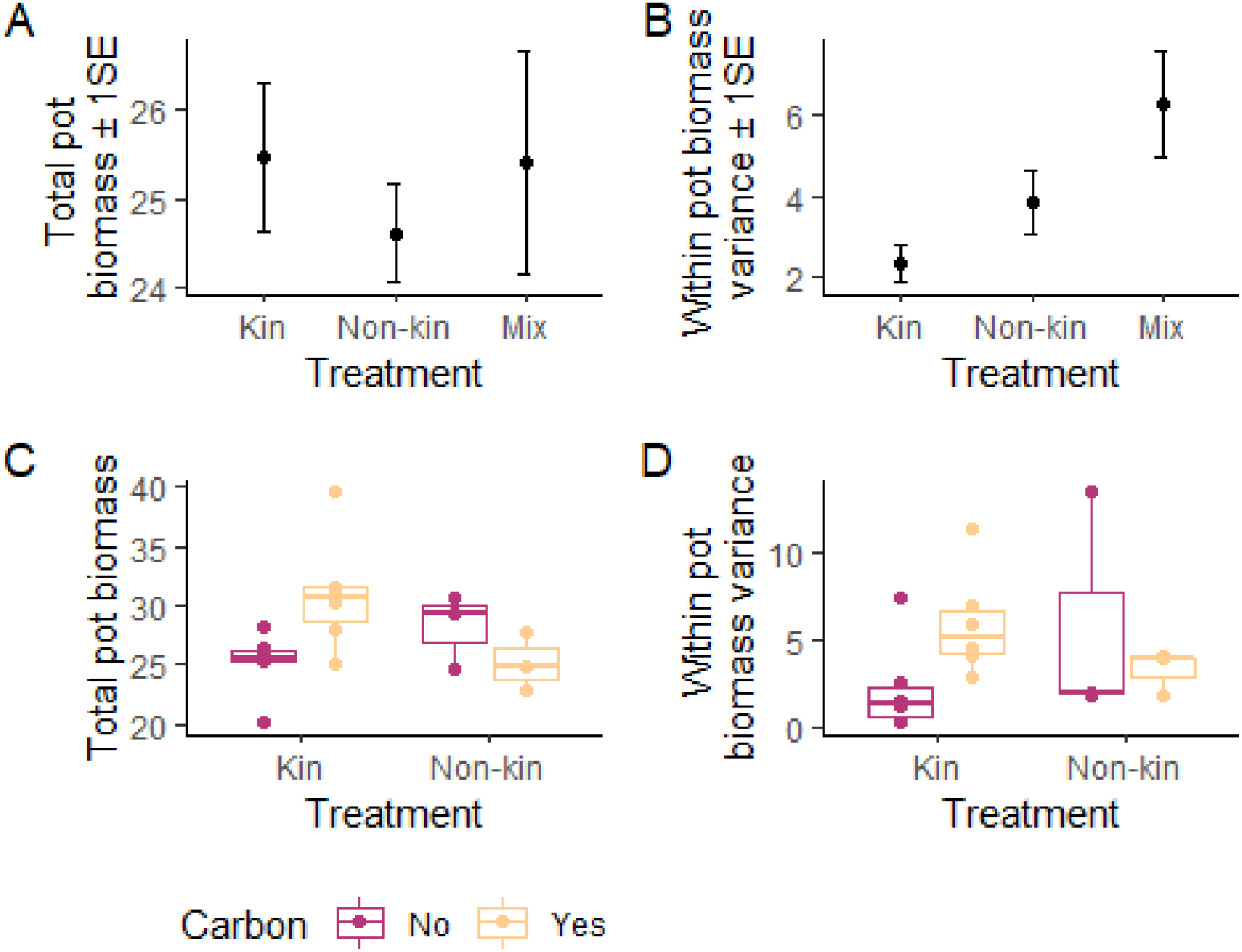
A, B) Plots representing mean total biomass (±1SE) and mean within-pot variance in biomass (±1SE) for untreated soil; C, D) Boxplots comparing total pot biomass and within-pot variance in biomass for kin and non-kin treatments in untreated soil *vs.* soil with activate carbon (experiment 2). Dots indicate single pots.

The differences among treatments in within-pot variance changed remarkably in pots treated with activated carbon. Total biomass in pots treated with activated carbon was higher compared to untreated pots (Table 4, Figure 4C). However, this effect was observed only in kin treatments as indicated by the significant treatment x carbon interaction. Similarly, activated carbon was associated with a large shift in within-pot variance in biomass for kin treatments (Table 4, Figure 4D). In kin treatments, within-pot variance in biomass for pots treated with activated carbon was on average 2.7 times larger than in untreated kin pots, and comparable to within-pot variance in biomass found in mix treatments in untreated soil (Figure 4B, D).

**Table 4:**
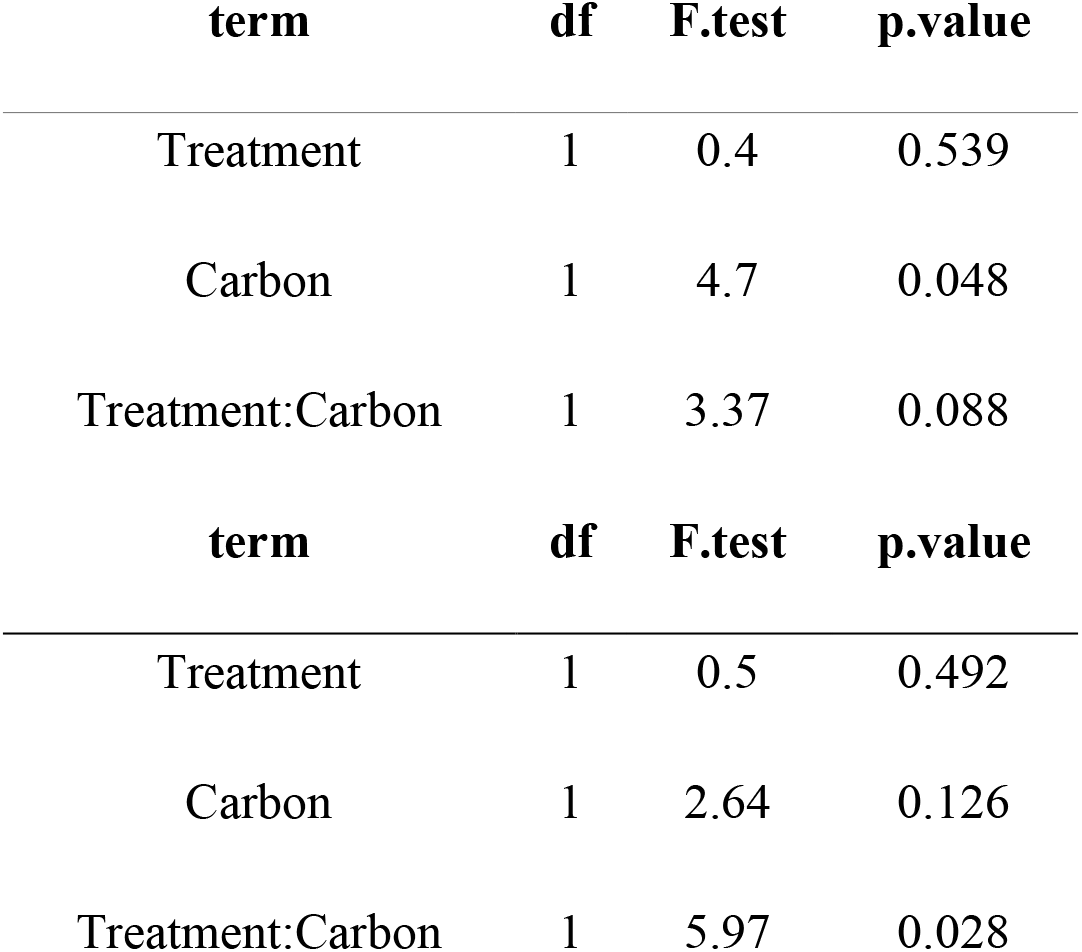
results of linear models on within pot biomass variance and total pot biomass (Experiment 2) in response to treatment (kin *vs.* non-kin), activated carbon application, and their interaction (N = 18 pots).

| term | df | F.test | p.value |
| --- | --- | --- | --- |
| Treatment | 1 | 0.4 | 0.539 |
| Carbon | 1 | 4.7 | 0.048 |
| Treatment:Carbon | 1 | 3.37 | 0.088 |
| term | df | F.test | p.value |
| Treatment | 1 | 0.5 | 0.492 |
| Carbon | 1 | 2.64 | 0.126 |
| Treatment:Carbon | 1 | 5.97 | 0.028 |

We found no effects of focal genotype identity (F_3_ = 2.31, p = 0.08) or treatment (F_2_ = 0.17, p = 0.83) on aboveground biomass of focal plants. However, to understand if biomass of focal plants varied with genotype identity of surrounding plants, we analysed the effect of surrounding and focal genotype identity on biomass of focal plants, in kin and non-kin treatments. This showed a significant effect of identity of surrounding genotype (F_3_ = 2.93, p = 0.04), but not of focal genotype (F_3_ = 0.59, p = 0.62). Focal plants surrounded by genotype 102 attained the highest biomass and focal plant surrounded by genotype 121 the lowest (Figure 5).

**Figure 5:**
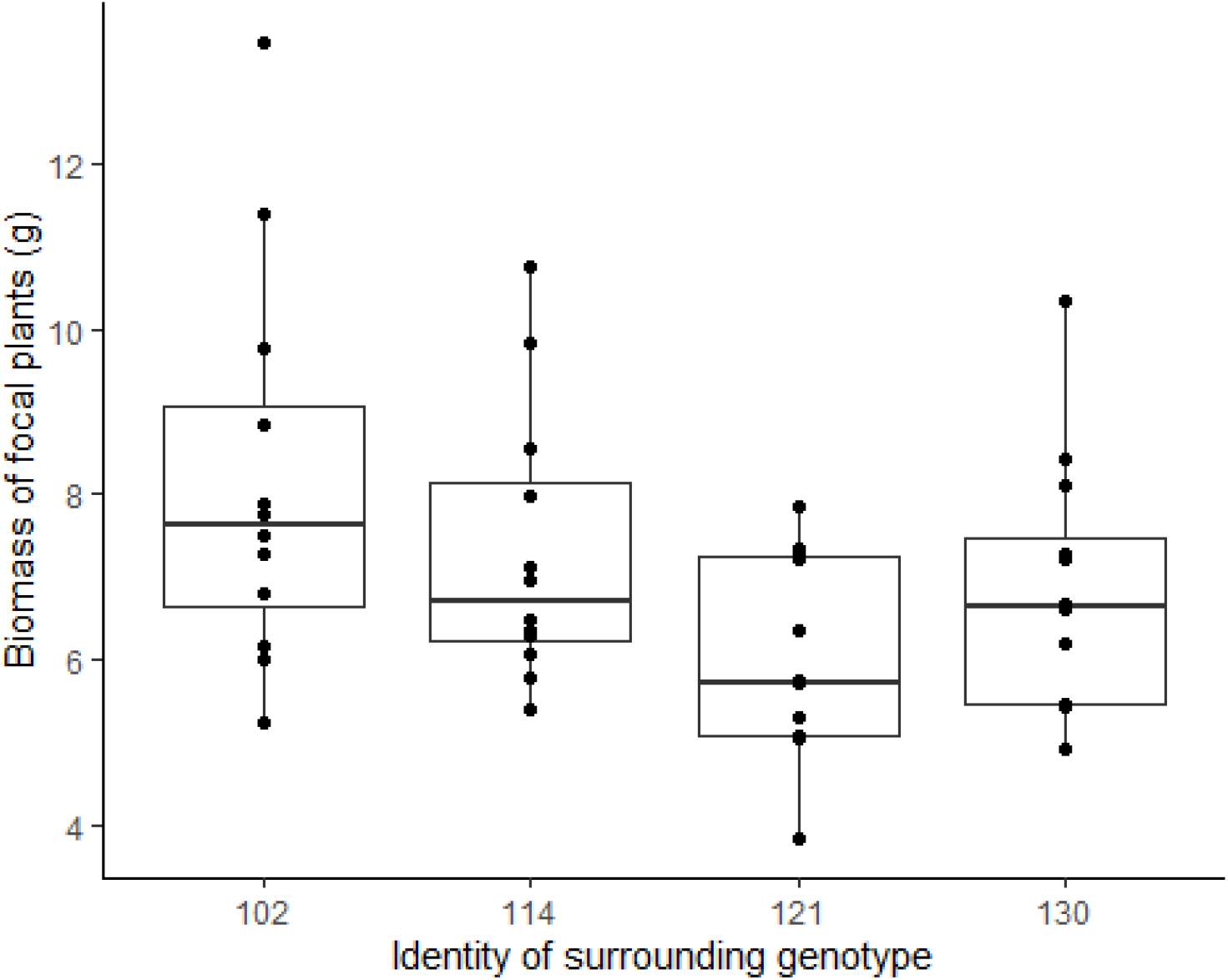
Boxplot representing biomass of focal plants in response to the identity of surrounding genotypes in kin and non-kin treatments (experiment 2). Dots represent single focal plants.

## Discussion

The results of our experiments provide support for both reduced competition among kin and genetic variation in competitive ability. Importantly, had we not specifically observed root behaviour in experiment 1, and added activated carbon to pots in the mini-communities of experiment 2, we would not have been able to conclude that kin recognition alters competitive growth.

Here we discuss separately the results pointing toward either mechanism (i.e. reduced competition among kin and genetic variation for competitive ability), and how these results may help better understand the different outcomes of intraspecific interactions. Our plants did not produce flowers before the end of either experiment. Hence, we were not able to test for fitness differences in seed set among genotypes, and we based our interpretations on the observed variations in root growth behaviour, biomass and growth among interacting plants.

### Reduced competition among kin

Our first experiment tested whether plants of *Medicago minima* altered their root growth toward their neighbour depending on whether the neighbour was kin or non-kin. We found that the response to neighbour relatedness was genotype-specific (Figure 1). In general, all genotypes grew more roots away from than toward their neighbours, which can be attributed to our experimental design, where two plants were grown in close proximity to each other leaving most space for root growth in the direction away from the neighbour.

Two of the focal genotypes showed a response consistent with avoiding kin root competition. Genotype 121 and genotype 114 showed differential growth response to kin *vs.* non-kin, and grew proportionally less roots toward their kin neighbours. In contrast, genotype 102 grew more roots toward its kin neighbour relative to non-kin. This genotype was also characterised by the slowest growth rate (Figure 2), and it is possible that for this genotype, growing with a kin of similar slow growth rate, simply resulted in more space to proliferate roots in direction of kin relative to fast-growing non-kin neighbours. Our findings of reduced root growth toward kin neighbours are consistent with previous studies also finding reduced root competition among kin (Dudley & File, 2007; Bhatt et al., 2010). It has been suggested that mechanisms generating kin *vs.* non-kin responses could be a side effect of self-avoidance (Yamawo, Sato, & Mukai, 2017; Novoplansky, 2019). However, irrespective of mechanisms, our observations on root behaviour point to a response to neighbour identity compatible with kin avoidance, and combined with the results from activated carbon treatments in our second experiment, suggest that kin avoidance is mediated by kin recognition.

In our second experiment, we exposed focal genotypes located at the centre of a pot, to three neighbours that where either kin, non-kin or a mix of stranger different genotypes. We found a positive relationship in early leaves growth of focal and surrounding plants in kin treatments, but not in non-kin and mix treatments (Figure 3 A-C). This response could be the result of reduced competition among kin, but it could also be explained by low variation in competitive ability among identical genotypes, leading to similar growth rates of focal and surrounding plants in kin treatments, and to weak or even negative relationships in growth rates of plants in non-kin and mixed treatments, where fast-growing and potentially more competitive genotypes could impede growth of slow-growing ones.

Further support for reduced competition among kin came from the results of total pot biomass and within-pot variance in biomass. We found no differences in total pot biomass among treatments (Figure 4A), indicating that regardless of treatment, plants grew to fill all the space and resource available within pots. However, we found large differences in how biomass was distributed among individual plants within the same pot (i.e. within-pot variance in biomass) depending on treatment (Figure 4B). This within-pot variance was the smallest in kin treatments and the largest in mix treatments, again suggesting reduced or very similar competition among plants in kin pots.

Application of activated carbon is known to inhibit the action of root exudates (Callaway & Aschehoug, 2000; Callaway et al., 2005), which have been demonstrated to mediate neighbour identity recognition in plants (Yang, Li, Xu, & Kong, 2018). Although the number of experimental units exposed to activated carbon treatments was low, the pattern we observed when comparing treated and untreated soil was striking. In kin treatments where activated carbon was added, within-pot biomass variance increase by almost 3 fold. Such changes were not observed in non-kin pots (Figure 4C, D).

The within-pot variance in biomass observed for kin treatments in activated carbon was comparable within-pot variance of mix treatments in untreated soil. This strongly suggests that when plants cannot detect the identity of their kin neighbours, they become more competitive. Reduced variance in biomass among plants in untreated kin pots is therefore most likely caused by kin recognition and reduced competition among kin. If this was a mere consequence of similarity in competitive ability among identical genotypes, we would not except such a strong shift in the variance of biomass among plants biomass in kin pots treated with activated carbon.

Although the reliability of activated carbon has been criticized as it may stimulate plants growth (Lau et al., 2008), our results are in line with previous studies (e.g. Biedrzycki, Jilany, Dudley, & Bais, 2010; Semchenko et al., 2014), which show that suppression of communication via root-exudates affects plant competition and alters the response to neighbour identity.

### Genetic variation in competitive ability

Our results also show presence of genotypic variation in competitive ability. For early radial growth of plants (Figure 3D), genotype 121 showed the fastest and genotype 102 the slowest growth rates. Consistent with this, we found that biomass of focal plants varied significantly depending on genotype identity of the surrounding plants (Figure 5). Focal plants attained the highest biomass when surrounded by genotype 102 (the genotype with the slowest growth), and the lowest biomass when surrounded by genotype 121 (the fastest growing genotype). This is consistent with other studies reporting intraspecific variation for competitive ability via variation in growth rate and ability to suppress growth of neighbours (e.g. Masclaux et al., 2010; Cahill, Kembel, & Gustafson, 2005).

Additional support for genotypic variation for competitive ability came from the significantly higher variance in biomass among plants within pot in mix treatments. This suggests a high asymmetric competition among different genotypes, consistent with genetic variation in competitive ability (Donohue, 2003; Simonsen et al., 2014).

### Intraspecific interactions as a result of both reduced kin competition and variation in competitive ability

Our results indicated that reduced competition among kin was most evident in genotypes characterized by the highest competitive ability, for which we considered growth and biomass to be a proxy. Although, the limited number of genotypes used for this experiment does not allow for broad generalizations, our results suggest that reduced competition among kin may not be independent of competitive ability. Across both experiments, genotype 102 showed the weakest response in terms of reduced competition towards kin, while attaining the lowest values of belowground biomass (experiment 1) and the smallest competitive effect on non-kin focal plants (experiment 2).

In contrast, genotype 121, which showed strong signals of reduced root competition among kin, showed the highest growth rate, obtained the larger biomass (experiment 1), and exerted the strongest competitive effect on focal plants (experiment 2). Similarly, genotype 114 also showed signals for reduced competition in kin treatments and was among the fastest growing and more competitive genotypes (experiment 1). This suggests that reduced competition among kin could be more pronounced in strongly competitive genotypes that are also more likely to interfere with their kin neighbours. Small genotypes may deplete resources more slowly than larger genotypes, thus the effects of competition with neighbours that may prompt a shift in root behaviour, would manifest comparatively later. This is consistent with previous studies showing that kin recognition is a density-dependent response (Lepik et al., 2012). Our experiment on root behaviour was terminated when roots of several plants reached the bottom of the pot and started growing horizontally. At this time, the amount of interference received from neighbouring roots may have been comparatively low for small genotypes (such as 102) exposed to equally small genotypes. This may also be a reason why we found no indication of reduced competition among kin in genotype 102. Our study is one of few (see also Biernaskie, 2010) showing experimentally that reduced kin competition and genetic variation in competitive ability act simultaneously to shape the outcome of intraspecific interactions.

In plant populations characterized by a strong spatial genetic structure, kin recognition and reduced kin competition may favour the growth of kin genotypes, creating positive density-dependent kin interactions. This can result in a higher resistance to invasion of more distantly related conspecifics and even of other species (Molofsky & Bever, 2002). Indeed, it was previously found that *M. minima* had higher survival in kin interactions compared to both non-kin and interspecific interactions (Ehlers et al., 2016). Positive density-dependent interactions in plants can influence both the distribution of intraspecific genetic variation and local species co-existence (Molofsky & Bever, 2002; Laird & Schamp, 2006). Moreover, it has been shown that such interactions can be important for invasion success and range expansion (Crandall & Knight, 2015).

Intraspecific interactions with more distantly related conspecifics should on the other hand be governed by variation in competitive ability among interacting genotypes (Fridley, Grime, & Bilton, 2007; Fridley & Grime, 2010). If highly competitive genotypes that do not reduce competition with kin, they may ultimately be subject to negative density-dependence, when increasing success results in more encounters with highly competitive kin genotypes. Our finding of genotypes that are both highly competitive against non-kin, and reduce their competition towards kin suggests a potentially highly successful strategy, unless curbed by trade-offs and costs of plastic growth response to neighbour’s identity (Crepy & Casal, 2014).

In conclusion, we found that within a species, intraspecific interactions are shaped by both reduced competition with kin and by variation in growth rate in response to non-kin neighbours. Had we not observed root behaviour, and suppressed neighbour recognition by adding activated carbon, the specific kin response might have gone unnoticed as the symmetric growth in kin pots could also be explained by genetic variation in competitive ability. When predicting outcome of intraspecific interactions, these findings emphasize the importance of knowing the degree of genetic substructure in populations as local positive density-dependent interactions may be expected in clusters of kin, but not of non-kin.

## Acknowledgments

This work was funded by a grant from the Danish Council of Independent Research (grant number 6108-00200B) to BKE. We are grateful to L. Lauridsen and J. Rytter for their help in the greenhouse.

## Authors contribution

BKE, MJ and CT planned the experiments, MJ and CT carried out the experiments, ST carried out the statistical analyses and wrote the first draft of the manuscript, all authors contributed to reviewing following versions.

## Data and code availability

Data is available on a Zenodo repository doi: 10.5281/zenodo.3441913. Statistical code is supplied in Appendix A2.

## Supplementary material

**Appendix 1**: **Figure A1**: **Photo taken while setting up experiment 1. Seedlings’ pairs were sowed at a distance of 2.5 cm, pins of different colours marked focal and neighbour genotypes. Containers (30 cm x 25 cm x 3 cm) were kept at a 50° angle, thus forcing roots to grow against the downward-facing surface and allowing their observation and measurement. To avoid exposure of roots to light, we kept the downward-facing side of each container covered with a black plastic slate that could be removed when recording root growth. Figure A2: Scans of root measurements in experiment 1. Different colours represent different measurement events. Lateral roots were classified as growing away or toward a neighbour. Plants were harvested once some of the individuals’ roots reached the bottom of the containers**.

**Appendix 2: Code used for data wrangling, statistical analyses and figures**.

