## Appendix 1 - Figure A1, A2 for "Intraspecific interactions in the annual legume *Medicago minima* are shaped by both genetic variation for competitive ability and reduced competition among kin"

**Figure A1: Photo taken while setting up experiment 1. Seedlings' pairs were sowed at a distance of 2.5 cm, pins of different colours marked focal and neighbour genotypes. Containers (30 cm x 25 cm x 3 cm) were kept at a 50° angle, thus forcing roots to grow against the downward-facing surface and allowing their observation and measurement. To avoid exposure of roots to light, we kept the downward-facing side of each container covered with a black plastic slate that could be removed when recording root growth.**

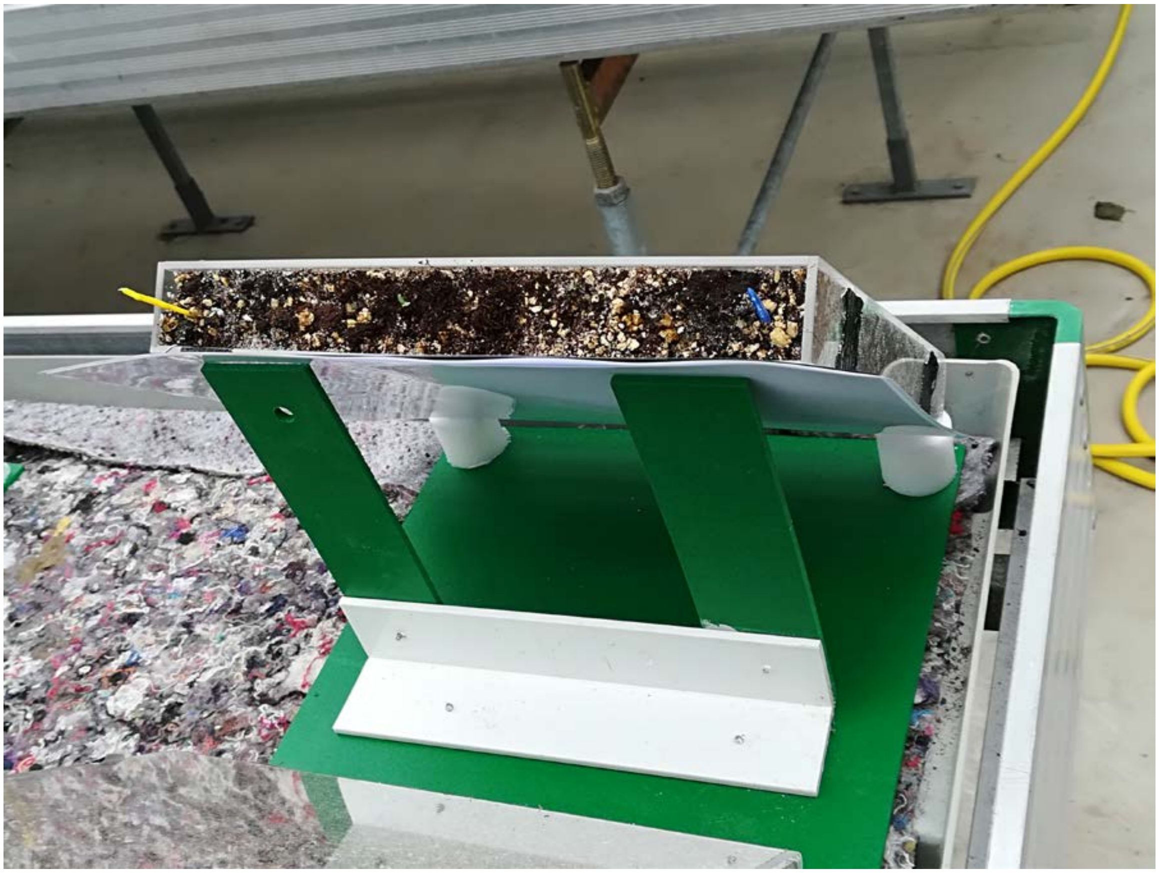

**Figure A2: Scans of root measurements in experiment 1. Different colours represent different measurement events. Lateral roots were classified as growing away or toward a neighbour. Plants were harvested once some of the individuals' roots reached the bottom of the containers.**

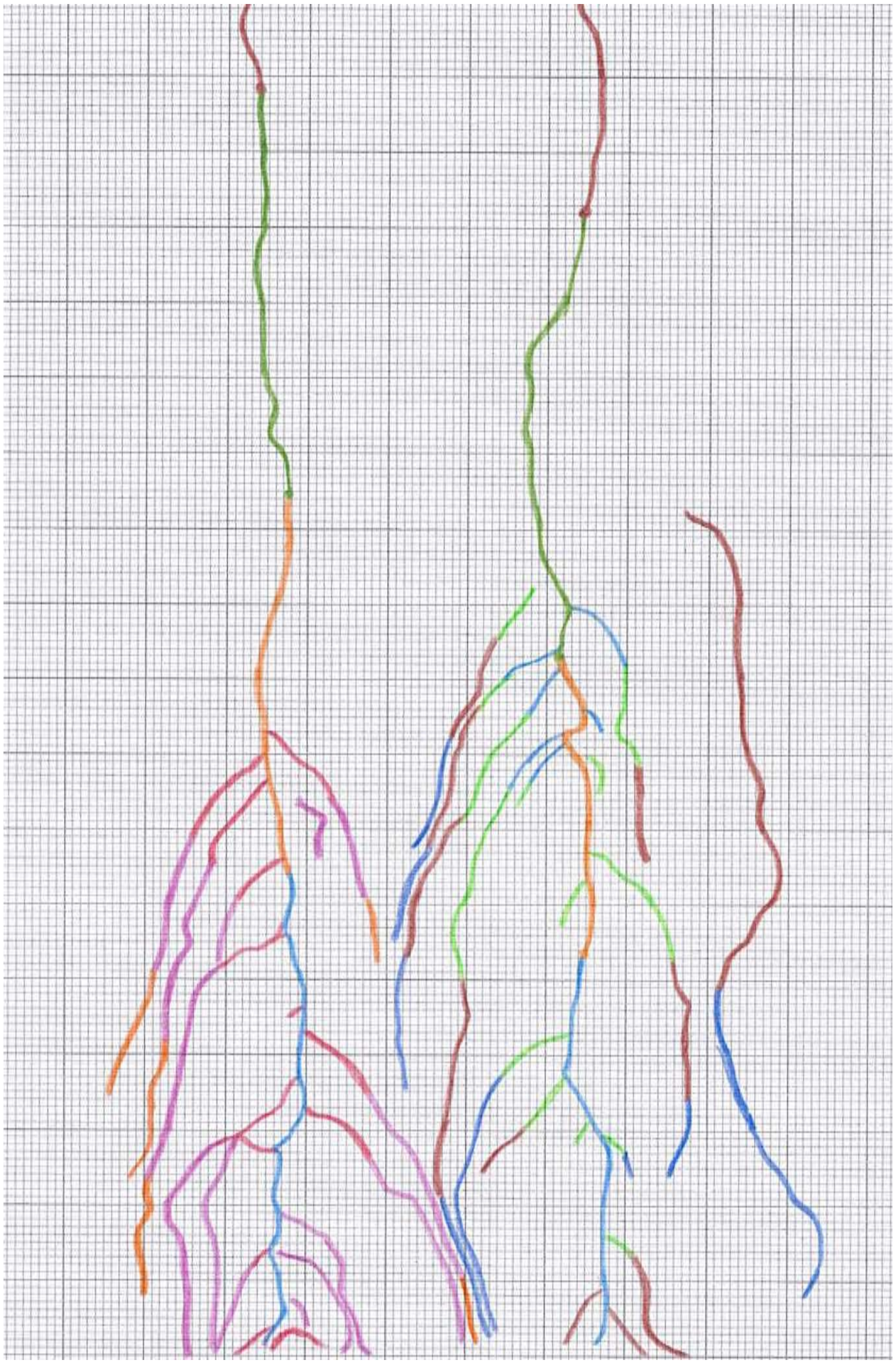

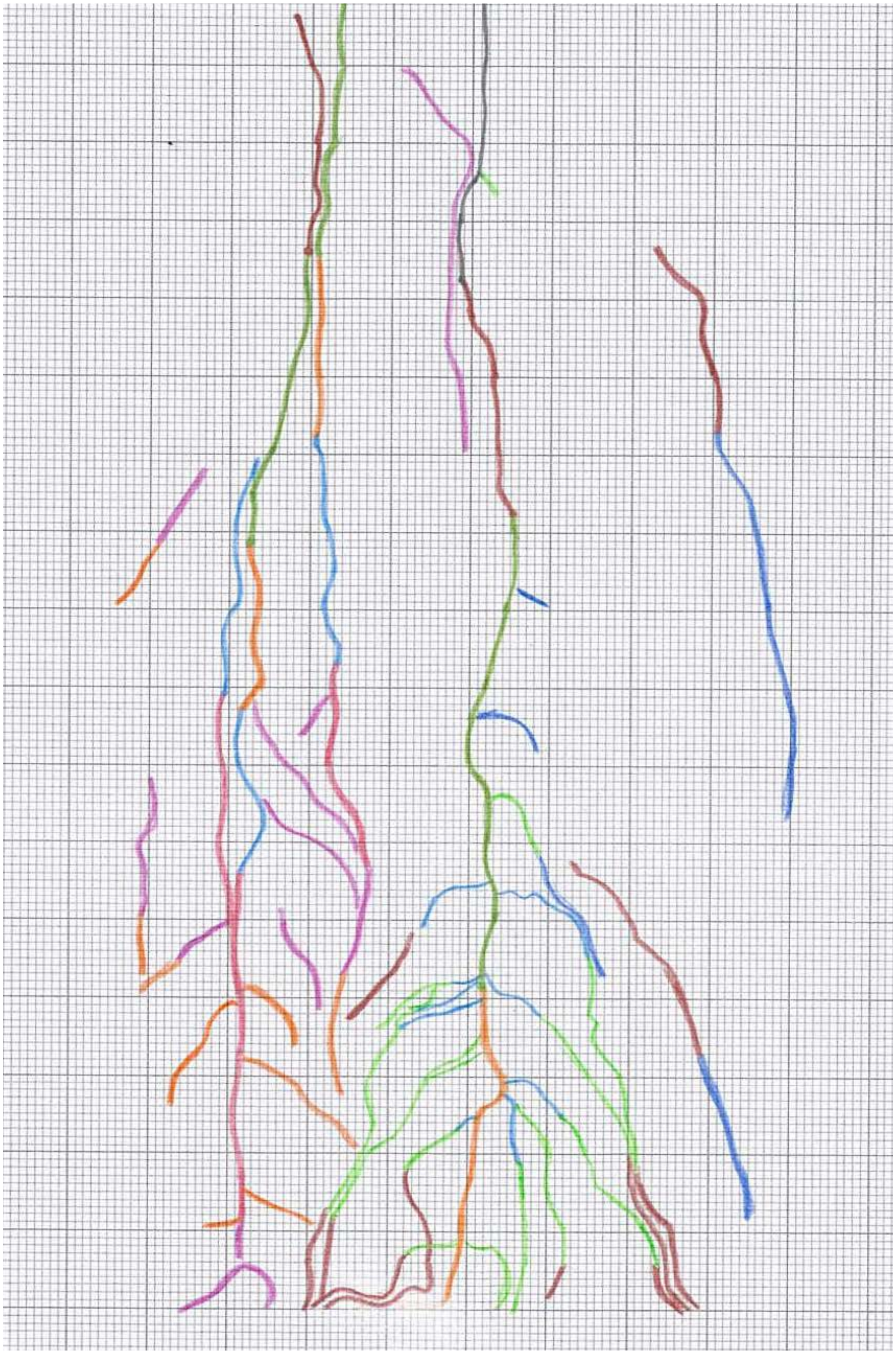
