## Appendix 2 for "Intraspecific interactions in the annual legume *Medicago minima* are shaped by both genetic variation for competitive ability and reduced competition among kin"

### Analyses Tomiolo et al Medicago minima

Setting up document and calling for R files containing data wrangling and list of dependencies and packages.

```
knitr::opts_chunk$set(echo = TRUE)
```

#### Data Preparation

Calling relevant libraries

```
library(tidyverse)
```

```
## -- Attaching packages -----------------------------------------------------------------------------
```

```
## v ggplot2 3.2.0     v purrr   0.3.2
## v tibble  2.1.3     v dplyr   0.8.2
## v tidyr   0.8.3     v stringr 1.4.0
## v readr   1.3.1     v forcats 0.4.0
```

```
## -- Conflicts ----------------------------------------------------------------------------------- tid
## x dplyr::filter() masks stats::filter()
## x dplyr::lag()    masks stats::lag()
```

```
library(patchwork)
library(viridis)
```

```
## Loading required package: viridisLite
```

```
library(ggpubr)
```

```
## Loading required package: magrittr
```

```
##
## Attaching package: 'magrittr'
```

```
## The following object is masked from 'package:purrr':
##
##     set_names
```

```
## The following object is masked from 'package:tidyr':
##
##     extract
```

```
library(pander)
library(emmeans)
library(pwr)
```

#### Import data files for analysis:

*Experiment 1: Root behaviour*

```
roots <- read_csv(here::here("data/root_behavior_experiment_for_ms.csv"))
```

```
## Parsed with column specification:
## cols(
##   .default = col_double(),
##   treatment_code = col_character(),
##   treatment = col_character()
## )

## See spec(...) for full column specifications.
```

*Experiment 2: Minicommunities*

```
minima_data <- read.csv(here::here("data/Medicago_minima_biomass_dataMerge.csv"))
```

read genotypes identifiers as factors

```
minima_data$Genotype_Focal <- as.factor(minima_data$Genotype_Focal)
minima_data$Genotype_Surround <- as.factor(minima_data$Genotype_Surround)
```

#### 1. Wrangling for root behaviour experiment

The original data file contains incremental measurements of root numbers and growth for each sampling date. To get the total number of roots and root growth we sum all the measurements together:

```
roots <- roots %>%
    mutate(total_root_toward  = rowSums(select(.,starts_with("root_toward"))),
           total_root_away = rowSums(select(., starts_with("root_away"))),
           total_growth_toward = rowSums(select(., starts_with("growth_toward"))),
           total_growth_away = rowSums(select(., starts_with("growth_away"))))
```

###### Proportion of roots number and of root growth growing away vs toward the neighbour, total biomass and root to total

```
roots <- roots %>%
    mutate(
        relative_root =
            (total_root_away - total_root_toward) /
            (total_root_away + total_root_toward),
        relative_root_growth =
            (total_growth_away - total_growth_toward) /
            (total_growth_away + total_growth_toward),
        total_biomass = abv_biomass + below_biomass,
        root_to_shoot = below_biomass / abv_biomass,
        root_to_total = below_biomass / (below_biomass + abv_biomass)
    )
```

Recoding non kin treatment initially named as "intra" in dataset

```
roots <- roots %>%
    mutate(treatment = fct_recode(treatment, "non kin" = "intra"))
```

Making sure that focal and neighbour genotype id are read as factors

```
roots$focal <- as.factor(roots$focal)
roots$neighbor <- as.factor(roots$neighbor)
```

#### 2. Wrangling for minicommunities experiment

Calculate early leaves and radial growth and total radial growth from the initial database

```
minima_data <- minima_data %>%
    dplyr::select(Treatment,
                  Treatment_Genotype,
                  Carbon,
                  PotNumber,
                  Replicate,
                  PlantPosition,
                  Genotype_Focal,
                  Genotype_Surround,
                  AbvBiomass,
                  NbOfLeaves1,
                  NbOfLeaves2,
                  NbOfLeaves3,
                  RadiusMax1,
                  RadiusMax2,
                  RadiusMax3,
                  RadiusMax4,
                  RadiusMax5,
                  RadiusMax6,
                  HeighMax1,
                  HeighMax2,
                  RadiusMax3) %>%
    mutate(Leaves_growth = (NbOfLeaves2 - NbOfLeaves1),
           RadiusGrowth = (RadiusMax2 - RadiusMax1),
           HeightGrowth = (HeighMax2 - HeighMax1),
           TotalRadiusGrowth = (RadiusMax6 - RadiusMax1))
```

##### Comparisons between focals and mean neighbours

Preparing files for testing whether growth of focal and mean growth of surrounding plants are related based on treatment (kin, non-kin and mix)

First we exclude pots treated with Carbon from the database and we select surrounding plants, then we calculate the mean growth (leaves, height and radius) and mean biomass of the three surrounding plants in each pot

```
Surroundings <-  minima_data %>%
    filter(PlantPosition == "Surrounding", Carbon == "No") %>%
    arrange(
        Treatment,
        PotNumber,
        PlantPosition,
        Genotype_Focal,
        Genotype_Surround,
        NbOfLeaves1,
        NbOfLeaves2
    ) %>%
    group_by(Treatment,
             PotNumber,
             PlantPosition,
             Genotype_Focal,
             Genotype_Surround) %>%
    summarize(
        mean_leaves = mean(Leaves_growth, na.rm = TRUE),
        mean_biomass = mean(AbvBiomass, na.rm = TRUE),
        mean_height = mean(HeightGrowth, na.rm = TRUE),
        mean_radius = mean(RadiusGrowth, na.rm = TRUE)
    ) %>%
    ungroup()
```

Then we apply the same for focal plants. Note that mean growth and biomass are the equivalent of the measured values, but need to be renamed consistently in order to join focals and surroundings in one file

```
Focals <-  minima_data %>%
    filter(PlantPosition == "Central", Carbon == "No") %>%
    arrange(
        Treatment,
        PotNumber,
        PlantPosition,
        Genotype_Focal,
        Genotype_Surround,
        NbOfLeaves1,
        NbOfLeaves2
    ) %>%
    group_by(Treatment,
             PotNumber,
             PlantPosition,
             Genotype_Focal,
             Genotype_Surround) %>%
    summarize(
        mean_leaves = Leaves_growth,
        mean_biomass = AbvBiomass,
        mean_height = HeightGrowth,
        mean_radius = RadiusGrowth
    ) %>%
    ungroup()
```

We then create a data frame for each response variable of interest

##### Leaves of focal and surrounding plants

We select the variables of interest from the Focals and Surrounding data frames and join them

```
Focal_leaves <- Focals %>%
    dplyr::select(Treatment,
                  PotNumber,
                  PlantPosition,
                  Genotype_Focal,
                  Genotype_Surround,
                  mean_leaves)
```

```
Surrounding_leaves <- Surroundings %>%
    dplyr::select(Treatment,
                  PotNumber,
                  PlantPosition,
                  Genotype_Focal,
                  Genotype_Surround,
                  mean_leaves)
```

```
minima_leaves <-
    bind_rows(Focal_leaves, Surrounding_leaves)
```

The resulting data frame has measurements of focal and surroundings in one column. We then create a data frame where focal and surroundings are on different columns

```
minima_leaves <- minima_leaves %>%
    spread(key = PlantPosition, value = mean_leaves)
```

##### Biomass of focal and surrounding plants

Same process followed for leaves measurements is applied here

```
Focal_biomass <- Focals %>%
    dplyr::select(
        Treatment,
        PotNumber,
        PlantPosition,
        Genotype_Focal,
        Genotype_Surround,
        mean_biomass
    )
```

```
Surrounding_biomass <- Surroundings %>%
    dplyr::select(
        Treatment,
        PotNumber,
        PlantPosition,
        Genotype_Focal,
        Genotype_Surround,
        mean_biomass
    )
```

```
minima_biomass <-
    bind_rows(Focal_biomass, Surrounding_biomass)
```

```
minima_biomass <- minima_biomass %>%
    spread(key = PlantPosition, value = mean_biomass)
```

###### Radial growth of focal and surrounding plants

Repeating the process applied for leaves and biomass

```
Focal_radius <- Focals %>%
    dplyr::select(Treatment,
                  PotNumber,
                  PlantPosition,
                  Genotype_Focal,
                  Genotype_Surround,
                  mean_radius)
```

```
Surrounding_radius <- Surroundings %>%
    dplyr::select(Treatment,
                  PotNumber,
                  PlantPosition,
                  Genotype_Focal,
                  Genotype_Surround,
                  mean_radius)
```

```
minima_radius <-
    bind_rows(Focal_radius, Surrounding_radius)
```

```
minima_radius <- minima_radius %>%
    spread(key = PlantPosition, value = mean_radius)
```

##### Total biomass and variance of biomass within pots

Calculate total biomass and variance of biomass among individuals within pots and make sure that duplicate elements/rows are removed

```
biomass_data <- minima_data %>%
    arrange(
        PotNumber,
        Treatment,
        Replicate,
        Treatment_Genotype,
        Carbon,
        PlantPosition,
        Genotype_Focal,
        Genotype_Surround,
        AbvBiomass
    ) %>%
    group_by(PotNumber,
             Treatment,
```

```
              Carbon,
              Genotype_Focal,
              Genotype_Surround) %>%
    summarize(
        TotalBiomass = sum(AbvBiomass, na.rm = TRUE),
        VarBiomass = var(AbvBiomass, na.rm = TRUE),
        Treatment_Genotype = unique(Treatment_Genotype),
        Replicate = unique(Replicate)
    ) %>%
    ungroup()
```

Select pots with untreated soil (i.e. exclude carbon treatments)

```
biomass_data_no_carbon <- biomass_data %>%
    filter(Carbon == "No")
```

Selecting treatments that have been exposed to activated carbon and untreated soil, making sure that there is an equal number of replicates for both untreated and treated soil

```
pot_biomass_carbon_comparison <- biomass_data %>%
    filter(Treatment_Genotype %in% c("K102", "K130", "NK114-102"),
           Replicate %in% c(1, 2, 3))
```

### Analyses

#### Analyses on database from Experiment 1

##### 1- Standardized difference in number of roots growing away from vs. toward neigbhours in response to identity of focal genotype and treatment (i.e. kin or non-kin)

Select relevant columns from database

```
roots_relative <- roots %>%
    dplyr::select(pot_nr, focal, neighbor, treatment, relative_root)
```

Apply the model and produce type III anova table

```
Model_roots_interaction <- lm(relative_root ~ focal * treatment , data = roots_relative)

Model_roots_interaction_table <- broom::tidy(car::Anova(Model_roots_interaction,
                       type = c("III")))
```

Conduct posthoc tests using pairwise comparisons on the interaction treatment x focal genotype, and subsequently estract estimates

```
emm_root_rel <- emmeans(Model_roots_interaction, pairwise ~ treatment|focal)
```

```
emm_root_rel_estimates <- summary(emm_root_rel)$emmeans %>% as_tibble()
```

Create a tidy output for model results

```
Model_roots_table_tidy <- Model_roots_interaction_table %>%
     dplyr::select(term, df, statistic, p.value) %>%
     mutate(F.test = round(statistic, 2)) %>%
     mutate(p.value = format.pval(round(p.value,digits = 3), eps = 0.001)) %>%
     dplyr::select(term, df, F.test, p.value) %>%
    dplyr::filter(term %in% c("focal", "treatment", "focal:treatment")) %>%
    dplyr::mutate(term = stringr::str_replace(term, "focal", "genotype_focal"))
```

##### 2- Standardized difference in total lentgh (cm) of roots growing away from vs. toward neighbours in response to identity of focal genotype and treatment (i.e. kin or non-kin)

Select appropriate columns from the database

```
roots_growth <- roots %>%
    dplyr::select(pot_nr, focal, neighbor, treatment, relative_root_growth)
```

Apply the model and produce Type III anova table

```
Model_roots_growth <- lm(relative_root_growth ~ focal * treatment , data = roots_growth)

Model_roots_growth_table <- broom::tidy(car::Anova(Model_roots_growth, type = c("III")))
```

Conduct posthoc tests using pairwise comparisons on the interaction focal genotype x treatment, and subsequently extract estimates

```
emm_root_growth <- emmeans(Model_roots_growth, pairwise ~ treatment|focal)

emm_root_growth_estimates <- summary(emm_root_growth)$emmeans %>% as_tibble()
```

Create a tidy output for model results

```
Model_roots_growth_table_tidy <- Model_roots_growth_table %>%
    dplyr::select(term, df, statistic, p.value) %>%
    dplyr::mutate(F.test = round(statistic, 2)) %>%
    dplyr::mutate(p.value = format.pval(round(p.value,digits = 3), eps = 0.001)) %>%
    dplyr::select(term, df, F.test, p.value) %>%
    dplyr::filter(term %in% c("focal", "treatment", "focal:treatment")) %>%
    dplyr::mutate(term = stringr::str_replace(term, "focal", "genotype_focal"))
```

##### 3- Models testing whether aboveground and belowground biomass, and root-to-total biomass ratio vary across focal gentypes and treatments (i.e. kin vs. non-kin)

###### a. aboveground biomass

Apply the model and produce type III anova table

```
above_interaction_gamma <- glm(abv_biomass ~ focal + treatment ,
                               family = Gamma(link = "identity"),
                               data = roots)
```

```
above_table <- broom::tidy(
    car::Anova(above_interaction_gamma,
               type = c("III")),
    test.statistic = c("Wald"),
    error,
    error.estimate =
        c("pearson",
          "dispersion",
          "deviance")
)
```

produce tidy output of results

```
above_table_tidy <- above_table %>%
    dplyr::select(term, statistic,df,p.value) %>%
    mutate(Wald.Chi.square = round(statistic, 2)) %>%
    mutate(p.value = format.pval(round(p.value, digits = 3),eps = 0.001)) %>%
    dplyr::select(term, Wald.Chi.square , df, p.value) %>%
    dplyr::filter(term %in% c("focal", "treatment")) %>%
    dplyr::mutate(term = stringr::str_replace(term, "focal", "genotype_focal"))
```

##### b. belowground biomass

Apply the model and produce the type III anova table

```
below_interaction_gamma <- glm(below_biomass ~ focal + treatment,
                               family = Gamma(link = "identity"),
                               data = roots)


below_table <- broom::tidy(car::Anova(below_interaction_gamma,
                                      type = c("III")),
                           test.statistic = c( "Wald"),
                           error, error.estimate =
                               c("pearson",
                                 "dispersion",
                                 "deviance"))
```

conduct posthoc tests to evaluate differences among focal genotypes using pairwise comparisons

```
emm_below <- emmeans(below_interaction_gamma, pairwise ~ focal)
```

produce tidy output of model results

```
below_table_tidy <- below_table %>%
    dplyr::select(term, statistic,df,p.value) %>%
    mutate(Wald.Chi.square = round(statistic, 2)) %>%
    mutate(p.value = format.pval(round(p.value, digits = 3),eps = 0.001)) %>%
    dplyr::select(term, Wald.Chi.square , df, p.value) %>%
    dplyr::filter(term %in% c("focal", "treatment")) %>%
    dplyr::mutate(term = stringr::str_replace(term, "focal", "genotype_focal"))
```

##### c. root-to-total biomass ratio

Apply the model and produce type III anova table

```
root_to_tot_normal <- lm(root_to_total ~ focal + treatment , data = roots)

root_to_tot_normal_table <- broom::tidy(car::Anova(root_to_tot_normal,
                        type = c("III")),
                        test.statistic = c( "Wald"))
```

Produce tidy output of results

```
root_to_tot_normal_table_tidy <- root_to_tot_normal_table %>%
    dplyr::select(term, statistic,df,p.value) %>%
    mutate(F.test = round(statistic, 2)) %>%
    mutate(p.value = format.pval(round(p.value, digits = 3),eps = 0.001)) %>%
    dplyr::select(term, F.test , df, p.value) %>%
    dplyr::filter(term %in% c("focal", "treatment")) %>%
    dplyr::mutate(term = stringr::str_replace(term, "focal", "genotype_focal"))
```

##### Power analysis for proportion of root numbers

```
#power analysis

power_roots_R <- pwr.f2.test(u = 3, v = 24, f2 = 0.4981, sig.level = 0.05)
```

##### Power analysis for proportion of root growth

```
#power analysis


power_growth_R <- pwr.f2.test(u = 3, v = 24, f2 = 0.5427, sig.level = 0.05)
```

#### Analyses on database from Experiment 2

##### 1- Regression between number of leaves of focal and neighbours (mean values)

Apply the model and produce type III anova table

```
model_leaves <- lm(sqrt(Central) ~ Treatment * Surrounding, data = minima_leaves)

model_leaves_table <- broom::tidy(car::Anova(model_leaves, type = c("III")))
```

Apply regression to each treatment separately and produce anova table

```
model_leaves_split <- minima_leaves %>%
    nest(-Treatment) %>%
    mutate(model = map(data, ~ lm(sqrt(Central) ~ Surrounding, data = .x)),
           tidy_output = map(model, ~ broom::tidy(car::Anova(.x, type = c("III"))))) %>%
    select(-data, -model) %>%
    unnest()
```

##### 2- Linear models on early radial growth of focal genotypes as a function of focal genotype identity and treatment (kin, non-kin, mix)

Apply model and produce anova table

```
model_radius <- lm(Central ~ Genotype_Focal *Treatment, data = minima_radius)
model_radius_table <- broom::tidy(car::Anova(model_radius))
```

Produce tidy output of model results

```
model_radius_table_tidy <- model_radius_table %>%
    select(term, df, statistic, p.value) %>%
    mutate(F.test= round(statistic, 2)) %>%
    mutate(p.value = format.pval(round(p.value, digits = 3),eps = 0.001)) %>%
    select(term, df, F.test, p.value) %>%
    filter(term %in% c("Genotype_Focal", "Treatment", "Genotype_Focal:Treatment"))
```

##### test if there are effects of focal and surrounding genotypes identity on focal plants biomass

using a subset of the database that considers only kin and non-kin treatments

```
minima_biomass_no_mix <- minima_biomass %>%
    filter(Treatment %in% c("K", "NK"))
```

Apply model (with log transformation which produced slightly better distributed residual plot) and produce anova table

```
model_subset_log <-
    lm(log(Central) ~ Genotype_Focal + Genotype_Surround, data = minima_biomass_no_mix)

model_subset_table <- broom::tidy(car::Anova(model_subset_log))
```

Ppairwise comparisons to test differences among surrounding genotypes

```
emmeans_surround <- emmeans(model_subset_log, pairwise ~ Genotype_Surround)
```

##### Within pot variance in biomass and total pot biomass across treatments (kin, non-kin, mix)

###### Within pot variance in biomass

apply model and produce anova table

```r
model_pot_variance_biomass <-
    lm(log(VarBiomass) ~ Treatment,data = biomass_data_no_carbon)

model_pot_variance_biomass_table <-
    broom::tidy(car::Anova(model_pot_variance_biomass,
                           type = c("III")))
```

posthoc tests via pairwise comparisons among treatments

```r
emm_variance <- emmeans(model_pot_variance_biomass, pairwise ~ Treatment)
```

###### Total pot biomass

Apply models and produce anova table

```r
model_pot_total_biomass <-
    lm(log(TotalBiomass) ~ Treatment,data = biomass_data_no_carbon)

model_pot_total_biomass_table <-
    broom::tidy(car::Anova(model_pot_total_biomass,
                           type = c("III")))
```

###### Effect of activated carbon treatments on total pot biomass and variance of biomass within pots (considers only kin and non-kin treatments and a subset of genotypes)

Apply model and produce table of results

```r
pot_carbon_variance <- lm(log(VarBiomass) ~ Treatment * Carbon,
                          data = pot_biomass_carbon_comparison)

pot_carbon_variance_table <-
    broom::tidy(car::Anova(pot_carbon_variance,
                           Type = c("III"), vcov. = NULL))
```

posthoc tests using pairwise comparisons for the interaction Carbon x Treatment (i.e. kin vs. non-kin)

```r
emmeans_carbon_var <-
    emmeans(pot_carbon_variance,
            pairwise ~ Carbon|Treatment)
```

produce tidy output of results

```r
pot_carbon_variance_table_tidy <- pot_carbon_variance_table %>%
    dplyr::select(term, df, statistic, p.value) %>%
    dplyr::mutate(F.test = round(statistic, 2)) %>%
    dplyr::mutate(p.value = format.pval(round(p.value, digits = 3), eps = 0.001)) %>%
    dplyr::select(term, df, F.test, p.value) %>%
    dplyr::filter(term %in% c("Treatment", "Carbon", "Treatment:Carbon"))
```

Produce a summary table with the mean values of variance of biomass within pot and total pot biomass across treatments (kin, non-kin) and carbon presence

```r
summary_table <- pot_biomass_carbon_comparison %>%
    group_by(Treatment, Carbon) %>%
    summarize(mean_var = mean(VarBiomass),
              tots = mean(TotalBiomass))
```

###### Total biomass

Apply the model and produce anova table

```r
pot_carbon_total <- lm(sqrt(TotalBiomass) ~ Treatment * Carbon,
                       data = pot_biomass_carbon_comparison)

pot_carbon_total_table <-
    broom::tidy(car::Anova(pot_carbon_total,
                           Type = c("III"), vcov. = NULL))
```

Run posthoc tests using pairwise comparisons for the interaction between carbon and treatment (kin vs non-kin)

```r
emmeans_carbon_total <- emmeans(pot_carbon_total, pairwise ~ Carbon|Treatment)
```

```
## Use 'contrast(regrid(object), ...)' to obtain contrasts on the response scale
```

produce tidy output of the model results

```r
pot_carbon_total_table_tidy <- pot_carbon_total_table %>%
    dplyr::select(term, df, statistic, p.value) %>%
    dplyr::mutate(F.test = round(statistic, 2)) %>%
    dplyr::mutate(p.value = format.pval(round(p.value, digits = 3), eps = 0.001)) %>%
    dplyr::select(term, df, F.test, p.value) %>%
    dplyr::filter(term %in% c("Treatment", "Carbon", "Treatment:Carbon"))
```

### Figures

**Figure 1: Boxplots of A) Relative number of roots growing away *vs.* toward the neighbour, B)Relative length of roots growing away *vs.* toward the neighbour. (experiment 1) Each dot represents one individual replicate. Positive values indicate that higher number of roots and higher root growth (length measured in cm) were recorded away *vs.* toward the neighbour. Panels represent focal genotypes (102, 114, 121, 130) separately, and on each panel we report the results of post-hoc tests for the effect of treatment. (kin *vs.* non-kin).**

Providing p-values for the posthoc tests

```r
#posthoc test p-values for number of roots
posthoc_root_rel <- data_frame(
    p_value = paste("p =",c(0.01, 0.06, 0.03, 0.29)),
    treatment = c("kin", "kin", "kin", "kin"),
    relative_root = c(0.95, 0.95, 0.95, 0.95),
    focal = c(102, 114, 121, 130)
)
```

```
## Warning: `data_frame()` is deprecated, use `tibble()`.
## This warning is displayed once per session.
```

```
#posthoc test p-values for length of roots
posthoc_root_growth <- data_frame(
    p_value = paste("p =",c(0.001, 0.04, 0.03, 0.34)),
    treatment = c("kin", "kin", "kin", "kin"),
    relative_root_growth = c(0.95, 0.95, 0.95, 0.95),
    focal = c(102, 114, 121, 130)
)
```

setting up the plots assembling the graphs

```
 #plotting the data relative to standardized differences in number   #of roots
plot_root_relative <- roots %>%
    mutate(kin_fill = if_else(focal == neighbor,"1","0")) %>%
    ggplot(aes(x = treatment, y = relative_root, colour = treatment)) +
    geom_boxplot() +
    geom_point(position = position_dodge(width = 0.75)) +
    geom_text(data = posthoc_root_rel,
              aes(label = p_value),
              show.legend = FALSE,
              nudge_x = 0.1,
              colour = "black",
              size = 3) +
    theme_classic() +
    theme(strip.background = element_blank(),
          strip.placement = "left",
          strip.text = element_text(face = "bold"),
          legend.position = "none",
          legend.title = element_blank()) +
    geom_hline(yintercept = 0, linetype = "dashed") +
    scale_colour_viridis_d(begin = 0.7, end = 0.9, labels = c("non kin", "kin")) +
    labs(x = "Treatment", y = "Proportion of number of roots \ngrowing away vs toward") +
    facet_grid(rows = vars(focal), scales = "free_x", switch = "y")


#plotting the data relative to standardized differences in length   #of roots
plot_root_relative_growth <- roots %>%
    mutate(kin_fill = if_else(focal == neighbor,"1","0")) %>%
    ggplot(aes(x = treatment, y = relative_root_growth, colour = treatment)) +
    geom_boxplot() +
    geom_point(position = position_dodge(width = 0.75)) +
    geom_text(data = posthoc_root_growth,
              aes(label = p_value),
              show.legend = FALSE,
              nudge_x = 0.1,
              colour = "black",
              size = 3) +
    theme_classic() +
    theme(strip.background = element_blank(),
          strip.placement = "left",
          strip.text = element_text(face = "bold"),
          legend.position = "none",
```

```
        legend.title = element_blank()) +
    geom_hline(yintercept = 0, linetype = "dashed") +
    scale_colour_viridis_d(begin = 0.7, end = 0.9, labels = c("non kin", "kin")) +
    labs(x = "Treatment", y = "Proportion of root length away vs toward") +
    facet_grid(rows = vars(focal), scales = "free_x", switch = "y")

#assembling the plot in one panel
figure1_panel <- plot_root_relative + plot_root_relative_growth +
    plot_layout(ncol = 2) + plot_annotation(tag_levels = "A")
```

**Figure 2: Boxplots of A) aboveground biomass, B) belowground biomass, C) root-to-total biomass ratio for the focal genotypes (experiment 1). Dots represent single focal plants**

```
#Aboveground biomass
above_biomass_figure <- roots %>%
    ggplot(aes(x = focal, y = abv_biomass)) +
    geom_boxplot() +
    geom_point(position = position_dodge(width = 0.75)) +
    theme_classic() +
    theme(strip.background = element_blank(),
          strip.placement = "left",
          strip.text = element_text(face = "bold"),
          legend.position = "bottom",
          legend.title = element_blank()) +
    scale_colour_viridis_d(begin = 0.7, end = 0.9) +
    labs(y = "aboveground biomass (g)", x = "focal genotype")

#belowground biomass
below_biomass_figure <- roots %>%
    ggplot(aes(x = focal, y = below_biomass)) +
    geom_boxplot() +
    geom_point(position = position_dodge(width = 0.75)) +
    theme_classic() +
    theme(strip.background = element_blank(),
          strip.placement = "left",
          strip.text = element_text(face = "bold"),
          legend.position = "bottom",
          legend.title = element_blank()) +
    scale_colour_viridis_d(begin = 0.7, end = 0.9) +
    labs(y = "belowground biomass (g)", x = "focal genotype")

#root-to-total biomass ratio
root_to_total_figure <- roots %>%
    ggplot(aes(x = focal, y = root_to_total)) +
    geom_boxplot() +
    geom_point(position = position_dodge(width = 0.75)) +
    theme_classic() +
    theme(strip.background = element_blank(),
          strip.placement = "left",
          strip.text = element_text(face = "bold"),
          legend.position = "bottom",
          legend.title = element_blank()) +
```

```
    scale_colour_viridis_d(begin = 0.7, end = 0.9) +
    labs(y = "root-to-total biomass ratio", x = "focal genotype")

#assembling the figure panel
figure2_panel <-
    above_biomass_figure +
    below_biomass_figure +
    root_to_total_figure +
    plot_layout(ncol = 2) +
    plot_annotation(tag_levels = "A")
```

**Figure 3: Regression for number of leaves grown during the first two weeks (experiment 2) on focal and surrounding genotypes across treatments (kin, non-kin and mix).**

```
#scatterplot of surrounding and central leaves in Kin treatments
kin.correl <- minima_leaves %>%
    filter(Treatment == "K") %>%
    ggplot(aes(x = Surrounding, y = Central, fill = Treatment)) +
    geom_point() +
    geom_smooth(colour = "black",method = "lm") +
    scale_fill_viridis(discrete = TRUE, begin = 0.1) +
    theme_bw() +
    theme(panel.grid = element_blank(),
          legend.position = "none") +
    labs(x = "Leaves growth \nneighbours", y = "Leaves gowth \nfocal") +
    ggtitle("Kin")

#scatterplot of surrounding and central leaves in non-kin treatments
nonkin.correl <- minima_leaves %>%
    filter(Treatment == "NK") %>%
    ggplot(aes(x = Surrounding, y = Central, fill = Treatment)) +
    geom_point() +
    geom_smooth(colour = "black",method = "lm") +
    scale_fill_viridis(discrete = TRUE, begin = 0.5) +
    theme_bw() +
    theme(panel.grid = element_blank(),
          legend.position = "none") +
     labs(x = "Leaves growth \nneighbours", y = "Leaves growth \nfocal") +
    ggtitle("Non-Kin")

#scatterplot of surrounding and central leaves in mix treatments
mix.correl <- minima_leaves %>%
    filter(Treatment == "Mx") %>%
    ggplot(aes(x = Surrounding, y = Central, fill = Treatment)) +
    geom_point() +
    geom_smooth(colour = "black", method = "lm") +
    scale_fill_viridis(discrete = TRUE, begin = 0.97) +
    theme_bw() +
    theme(panel.grid = element_blank(),
          legend.position = "none") +
     labs(x = "Leaves growth \nneighbours", y = "Leaves growth \nfocal") +
    ggtitle("Mix")
```

```r
#early radial growth of focal genotypes
radial <- minima_radius %>%
    ggplot(aes(x = Genotype_Focal, y = Central)) +
    geom_boxplot() +
    geom_point(position = position_dodge(width = 0.75)) +
    theme_classic() +
    labs( x = "Focal genotype", y = "Radial growth")

#assembling the final panel
Figure3_panel <-
    kin.correl +
    nonkin.correl +
    mix.correl +
    radial +
    plot_layout(ncol = 2) +
    plot_annotation(tag_levels = "A")
```

**Figure 4: A,B) Plots representing mean total biomass (±1SE) and mean within-pot variance in biomass (±1SE) for untreated soil; C,D) Boxplots comparing total pot biomass and within-pot variance in biomass for kin and non-kin treatments in untreated soil vs. soil with activate carbon (experiment 2). Dots indicate single pots**

recoding the labels of treatments to assure consistency in graphics labels

```r
biomass_data_no_carbon <-
    biomass_data_no_carbon %>%
    mutate(
        Treatment = fct_recode(
            Treatment,
            "Non-kin" = "NK",
            "Kin" = "K",
            "Mix" = "Mx")) %>%
    mutate(
        Treatment = factor(
            Treatment,
            levels = c("Kin", "Non-kin", "Mix")))

pot_biomass_carbon_comparison <-
    pot_biomass_carbon_comparison %>%
    mutate(Treatment = fct_recode(
        Treatment,
        "Non-kin" = "NK" ,
        "Kin" = "K"))
```

calculating mean and standard errors for graphs

```r
#total biomass
total_sum <- biomass_data_no_carbon %>%
    group_by(Treatment) %>%
    summarize(mean_biomass = mean(TotalBiomass),
              error_biomass = sd(TotalBiomass)/sqrt(length(TotalBiomass))) %>%
    ungroup() %>%
```

```
    mutate(se_min = mean_biomass - error_biomass,
           se_max = mean_biomass + error_biomass)

#variance of biomass within pots
total_variance <- biomass_data_no_carbon %>%
    group_by(Treatment) %>%
    summarize(mean_variance = mean(VarBiomass),
              error_variance = sd(VarBiomass)/sqrt(length(VarBiomass))) %>%
    ungroup() %>%
    mutate(sev_min = mean_variance - error_variance,
           sev_max = mean_variance + error_variance)
```

plotting the graphs

```
#total biomass
total <- total_sum %>%
    ggplot(aes(x = Treatment, y = mean_biomass, ymin = se_min, ymax = se_max)) +
     geom_errorbar(position = position_dodge(width = 0.1), width = 0.1) +
    geom_point(position = position_dodge(width = 0.1), width = 1) +
    theme_classic() +
    labs( x = "Treatment", y = "Total pot \nbiomass \u00B1 1SE")
```

```
## Warning: Ignoring unknown parameters: width
```

```
#variance of biomass within pots
variance <- total_variance %>%
    ggplot(aes(x = Treatment, y = mean_variance, ymin = sev_min, ymax = sev_max)) +
     geom_errorbar(position = position_dodge(width = 0.1), width = 0.1) +
    geom_point(position = position_dodge(width = 0.1), width = 1) +
    theme_classic() +
    labs( x = "Treatment", y = "Within pot biomass \nvariance ± 1SE")
```

```
## Warning: Ignoring unknown parameters: width
```

```
#total biomass in activated carbon vs untreated pots
total_c <- pot_biomass_carbon_comparison %>%
    ggplot(aes(x = Treatment ,y = TotalBiomass, colour = Carbon)) +
    geom_boxplot() +
    geom_point(position = position_dodge(width = 0.75)) +
    theme_classic() +
     theme(legend.position = "bottom") +
    scale_colour_viridis_d(option = "A", begin = 0.5, end = 0.9) +
    labs( x = "Treatment", y = "Total pot biomass")
```

```
#variance of biomass within pots in activated carbon vs untreated pots
variance_c <- pot_biomass_carbon_comparison %>%
    ggplot(aes(x = Treatment ,y = VarBiomass, colour = Carbon)) +
    geom_boxplot() +
    geom_point(position = position_dodge(width = 0.75)) +
    theme_classic() +
    theme(legend.position = "none") +
    scale_colour_viridis_d(option = "A", begin = 0.5, end = 0.9) +
    labs( x = "Treatment", y = "Within pot \nbiomass variance")
```

Assembling the plots in the figure panel

```
Figure_4_panel <-
    total +
    variance +
    total_c +
    variance_c +
    plot_layout(ncol = 2) +
    plot_annotation(tag_level = "A")
```

**Figure 5: Boxplot representing biomass of focal plants in response to the identity of surrounding genotypes in kin and non-kin treatments (experiment 2). Dots represent single focal plants.**

```
Biomass_by_surrounding <-
    minima_biomass_no_mix %>%
    ggplot(aes( x = Genotype_Surround, y = Central)) +
    geom_boxplot() +
    geom_point(position = position_dodge(width = 0.75)) +
    theme_classic() +
    theme(panel.grid = element_blank()) +
    labs(x = "Identity of surrounding genotype", y = "Biomass of focal plants (g)")
```
